# Infinity Flow: High-throughput single-cell quantification of 100s of proteins using conventional flow cytometry and machine learning

**DOI:** 10.1101/2020.06.17.152926

**Authors:** Etienne Becht, Daniel Tolstrup, Charles-Antoine Dutertre, Florent Ginhoux, Evan W. Newell, Raphael Gottardo, Mark B. Headley

**Author notes:** These authors contributed equally (Evan W. Newell, Raphael Gottardo, Mark B. Headley).

## Abstract

Modern immunologic research increasingly requires high-dimensional analyses in order to understand the complex milieu of cell-types that comprise the tissue microenvironments of disease. To achieve this, we developed Infinity Flow combining hundreds of overlapping flow cytometry panels using machine learning to enable the simultaneous analysis of the co-expression patterns of 100s of surface-expressed proteins across millions of individual cells. In this study, we demonstrate that this approach allows the comprehensive analysis of the cellular constituency of the steady-state murine lung and to identify novel cellular heterogeneity in the lungs of melanoma metastasis bearing mice. We show that by using supervised machine learning, Infinity Flow enhances the accuracy and depth of clustering or dimensionality reduction algorithms. Infinity Flow is a highly scalable, low-cost and accessible solution to single cell proteomics in complex tissues.

## 1 Main

One of the cornerstones of immunology has been the detailed phenotypic classification of the cellular populations of the immune system. These efforts have largely been guided by identifying heterogeneity in surface protein expression across cell types. However, the breadth of this classification has been hindered by technical requirements allowing the analysis of only small subsets of markers per sample and analytical tools which are generally manual and low throughput. Over the past decade, methods that enable deeper interrogation of cellular heterogeneity in complex tissues and systems have provided a better understanding of the mechanistic underpinnings of disease. The key to this idea is being able to simultaneously measure large numbers of parameters on individual cells. This increased dimensionality facilitates the understanding of the unique characteristics of each individual cell and how cells interact within a given system. Modern flow cytometric approaches exemplify this by using panels of multiple fluorochrome-(conventional flow cytometry) or metal-(mass cytometry) conjugated antibodies to measure protein-expression profiles of individual cells with high cell throughput in order to capture and analyze both common and rare cell populations. With current instrumentation, fluorescence and mass cytometric approaches are unfortunately limited to 40 or fewer parameters. However, at least 371 cluster of differentiation (CD) markers are currently recognized (Engel et al. 2015), not including relevant surface proteins without CD designation. This disparity indicates that we are vastly limited in the comprehensiveness of these assays. In addition, robust implementation of panels approaching 40 parameters requires extensive experience and optimization as well as dedicated and uncommon instrumentation. Recently, several approaches have been published which utilize oligonucleotide-conjugated antibodies in combination with single cell RNA-sequencing technology to simultaneously measure single cell transcriptomes and 100s of surface proteins (Stoeckius et al. 2017; Hwang et al. 2020). Unfortunately, these techniques are still limited in terms of cellular throughput (10,000s of single cells), remain expensive (requiring deep sequencing and large antibody panels) and often require complicated experimental protocols. As such, there is a need for methods enabling concordant analysis of the expression patterns of high numbers of proteins at a single cell level while retaining both accessibility, low cost and high cell throughput.

Parallel to these experimental developments, computational techniques have also matured and found new applications in biology. Numerous studies have shown that independent single-cell experiments can be combined into a single augmented dataset, suggesting that multiple and partially-overlapping datasets can be leveraged to obtain a single integrated and informative data matrix (Abdelaal et al. 2019; Pedreira et al. 2008; Leite Pereira et al. 2019; Tinnevelt et al. 2019; Haghverdi et al. 2018; Stuart et al. 2019). Machine learning techniques have also significantly improved in the last decade, with well-known applications to fields such as genomics, computer vision or speech recognition (Eraslan et al. 2019;

LeCun, Bengio, and Hinton 2015). Although deep learning (using deep neural networks) is one of the most popular techniques at the moment, machine learning encompasses a large number of tools including regularized regression (Tibshirani 1996), support vector machines (Chang and Lin 2011) or ensemble of decision trees (Hastie et al. 2009; Breiman 2001; Chen and Guestrin 2016) to name a few. A common feature of these algorithms is their ability to model non-linear relationships. Non-linearity corresponds to relationship between variables that cannot be accurately modeled by a straight line (or its highdimensional equivalent). In the context of single cell analyses, the relationship between expression levels of a set of surface proteins is a case of a non-linear system. This non-linear system is often implicitly modeled as a Boolean formula, for instance “CD45 expression *and* CD19 expression *implies* CD79b expression”. Immunologists have extensively published such limited non-linear relationships between marker expression and cell phenotypes in the past decades. The actual system is however more subtle, as cell surface proteins are not categorical but continuous variables, depend on the biological context, and can be multimodal.

In this article, we demonstrate the capabilities of Infinity Flow, a low-cost methodology and computational toolset for high-dimensional single cell analysis of complex cell suspensions using standard flow cytometry instrumentation, off-the-shelf reagents, and scalable antibody panels. Infinity Flow uses machine learning to systematize imputation of cell surface protein expression at the single-cell level from experimental flow cytometry data. We leverage massively-parallel cytometry (MPC) experiments, which sparsely measure hundreds of exploratory antibodies and exhaustively measure a small number of well-characterized and informative so-called *‘‘Backbone”* antibodies. Machine learning is then used to impute expression levels of each exploratory marker for each single-cell based on non-linear functions of the measured *Backbone* markers. Infinity Flow enables the concurrent analysis of co-expression patterns of 100s of surface proteins across millions of cells at single cell resolution at a fraction of a cent per cell (USD). Importantly, this approach can be implemented on any existing flow cytometry platform with off-the-shelf reagents, rendering it highly accessible. We previously illustrated a proof-of-concept approach to delineate circulating conventional dendritic cells 2 (cDC2) and circulating monocytes (Dutertre et al. 2019), or to resolve the developmental trajectory of the neutrophil lineage during hematopoiesis (Kwok et al, in press). In this article, we present the complete and expanded methodology including an open-source R package *infinityFlow* submitted to the Bioconductor repository, optimized machine learning algorithms with benchmarks, single-cell background correction, and high-dimensional computational analyses of millions of single cells from the murine lung at steady-state and during metastatic seeding. Using our approach, we show that non-linear regression is superior to linear regression for machine learning based prediction of flow cytometric data. Further, we highlight the ability for Infinity Flow to augment a standard flow cytometric panel and enable more robust population and protein expression characterization. We use this approach to comprehensively define the cell populations of the murine lung, only using a 15 color flow cytometry panel. Finally, we identify two populations of tumor-ingesting macrophages during early metastatic seeding of the lung by melanoma cells that can be separated by PD-L1, MHCII and CD11c expression.

## 2 Results

### 2.1 The Infinity Flow pipeline

Infinity Flow processes data from low-cost (Supplementary Table 1) commercially available plate-based antibody screening panels (which we term massively-parallel cytometry, MPC), using machine learning to achieve the simultaneous analysis of hundreds of surface markers on millions of single cells isolated from complex tissues. Similar to conventional flow cytometry, MPC experiments start by staining a sample with a cocktail of fluorescently-labeled antibodies, here termed the *Backbone* panel (Figure 1a). Critical to the MPC approach, the *Backbone* panel should be designed with at least one empty fluorescence channel (usually PE or APC) and optimized to minimize spectral overlap into this open channel. The sample is then aliquoted across *w* wells (typically *w* ≈ 300) (Amir et al. 2019; Collier et al. 2017; Dutertre et al. 2019; Graessel et al. 2015; Koh et al. 2016; Uezumi et al. 2016; Kalina et al. 2019) (Figure 1b). Each well contains a distinct antibody clone conjugated to the fluorophore not used for the Backbone. We term the set of well-specific markers the *Infinity* panel. After completion of the staining step, each well contains a fraction of the total sample, and all cells within are stained with all Backbone markers in addition to a single *Infinity* marker (Figure 1c). Each well is then acquired as an individual flow cytometry sample using a conventional flow cytometer. Commercial kits exist from various vendors with a range of 240 - 371 individual antibodies, these kits can be further augmented with researcher-selected antibodies as needed.

**Figure 1.**
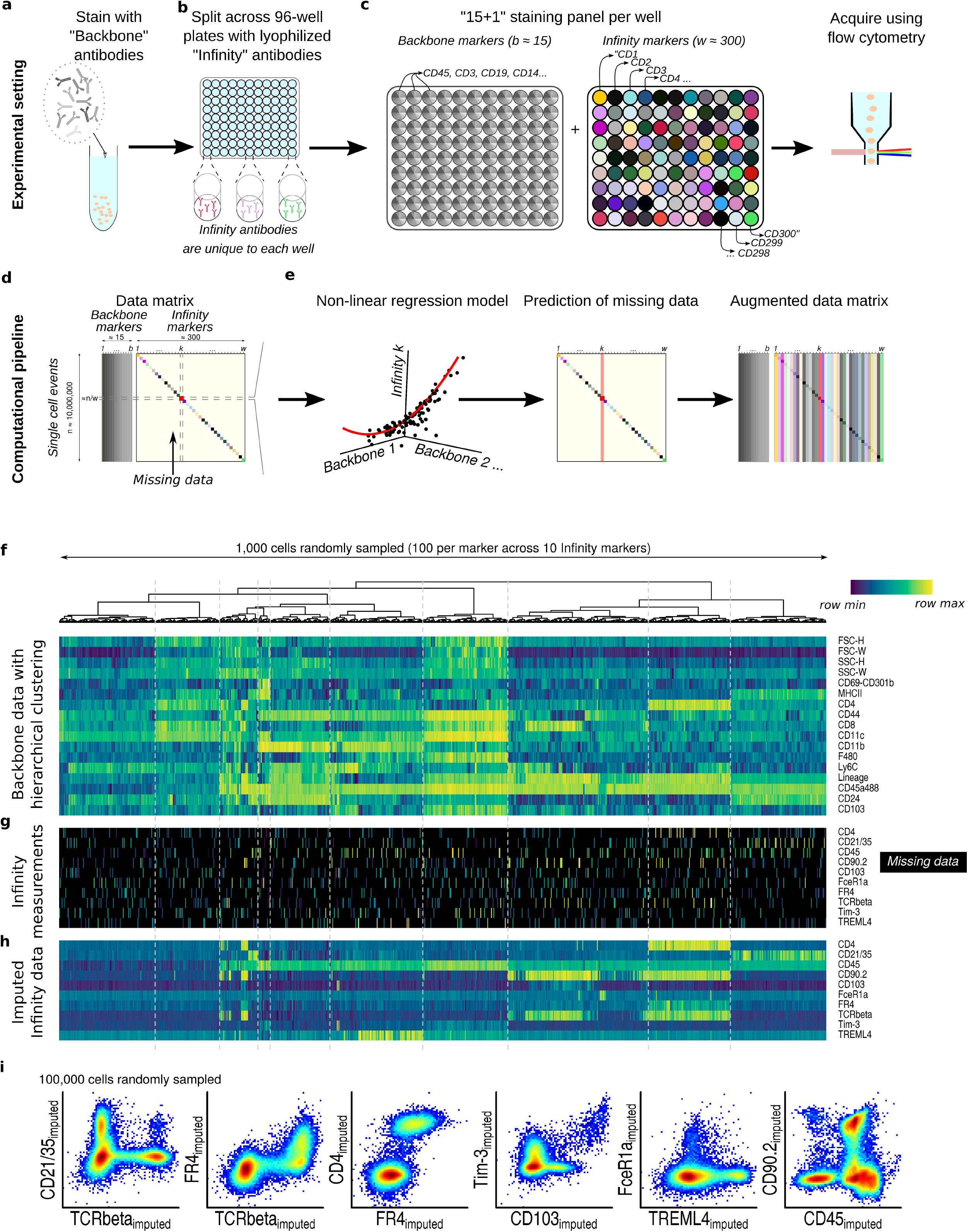
Summary of the Infinity Flow experimental design and computational pipeline. Experimental pipeline: **a)** Backbone panel staining, **b)** Infinity panel staining, **c)** Per-well staining panels and data acquisition. Computational pipeline: **d)** Data matrix with dense Backbone and sparsely non-missing Infinity markers measurements, **e)** Fitting of per-well non-linear regression models and missing data imputation. Example: **f)** on 1,000 cells, Backbone matrix with hierachical clustering of cells, **g)** its corresponding sparse Infinity markers measurements and **h)** its corresponding dense Infinity Flow imputed data. **i)** Imputation of co-expression patterns by Infinity Flow.

Each well-specific dataset (stored in its own FCS file), resulting from an MPC experiment, contains empirical measurements of every Backbone marker and empirical measurement of one of the Infinity panel markers. The data matrix resulting from all wells can be viewed as a dense matrix of *n* events ×*b* Backbone markers jointly measured with a sparse n ×w matrix of Infinity markers (Figure 1d). Following standard quality control of the data (Methods) the data is analyzed using non-linear multivariate regression to recast this disjointed data structure into a single cohesive expression matrix of all markers across all cells. To achieve this, we train, for every well, a machine learning model that predicts the expression of the Infinity panel marker on a continuous scale from the measured Backbone markers intensities. Once trained, these models are applied across the whole dataset to estimate the intensity of each of the w Infinity antibodies across the n events, resulting in a n ×w *dense* Infinity matrix of imputed intensities (Figure 1e).

This computational workflow is illustrated on cells isolated via collagenase digestion of non-perfused whole mouse lungs and stained using a standard 14-color immunoprofiling Backbone panel and Biolegend Murine LEGENDScreen MPC kit. The lung, even under homeostatic conditions, contains an exceptionally diverse cellular milieu, comprised of common and rare, immune and non-immune cells, providing an ideal testing ground for our high-dimensional approach. For this simplified example, we sub-sampled a total of 1,000 mouse lung cells from 10 wells each containing an antibody to a distinct CD molecule. We used hierarchical clustering to highlight structure within the Backbone (Figure 1f). More importantly, this structure correlates with distinct expression patterns in the sparse Infinity marker measurements (Figure 1g). The goal of Infinity Flow is to model these correlations in a data-driven manner using machine learning. These models then impute the expression of each Infinity panel marker across every cell in the dataset (Figure 1h). The dense, continuous and single cell data format of Infinity Flow’s output enables easy visualization and exploration of any combination of co-expression patterns across both the Backbone and Infinity panels (Figure 1i). Infinity Flow’s output is notably compatible with standard flow cytometry analysis software (e.g. FlowJo or flowCore (Hahne et al. 2009)), and can be used as input for further downstream analyses such as clustering, dimensionality reduction or automated gating.

### 2.2 Non-linear regression models accurately impute cytometry data

In order to maximize the quality of the predictions of the Infinity Flow algorithm, we quantitatively assessed its performance across Infinity markers and machine learning algorithms on the mouse lung cell dataset. Protein expression on cells is a continuous variable. However, standard flow cytometry analysis generally involves a progressive gating scheme whereby expression levels are discretized into two or more bins (e.g. low expression versus high expression), with potentially imbalanced frequencies (e.g. many negative events and few positive events). In order to account for such imbalance and pseudo-discrete format, we decided to use the Area under the ROC curve (AUC) on held-out data as a performance metric as opposed to continuous metrics such as the *R^2^* coefficient (Methods), focusing our effort on the magnitude and frequency of imputed expression.

The Infinity Flow R package supports a variety of linear and non-linear regression based machine learning models. Given the expected non-linear relationships of protein expressions, we hypothesized that non-linear models would outperform linear models in this setting. We therefore compared the performance of the non-linear algorithms (Chollet and Allaire 2018; Chen and Guestrin 2016; Chang and Lin 2011; Tibshirani 1996) to linear L1-penalized models (Tibshirani 1996). Consistent with our hypothesis, non-linear methods significantly outperformed linear models (Figure 2a). Overall, each nonlinear regression method performed well, with a median AUC between 0.88 and 0.89 (Supplementary Table 2), while the linear model performed much worse (median AUC of 0.71). Gradient boosted trees implemented by the XGBoost library (Chen and Guestrin 2016), performed slightly better in this benchmark (Figure 2a and Supplementary Figure 1).

**Figure 2.**
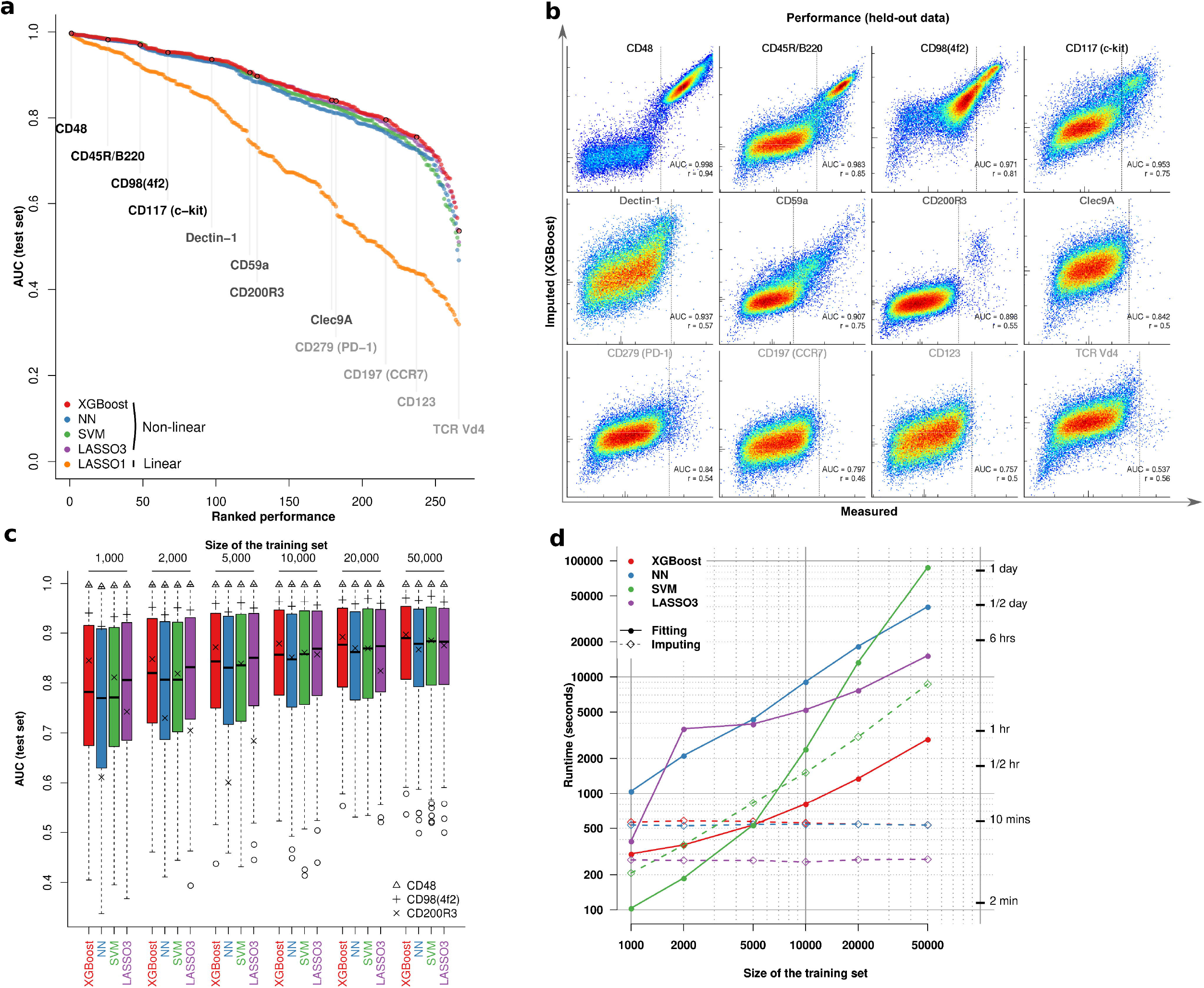
Non-linear regression models accurately impute cytometry data. **a)** AUC (computed using manual gating as ground truth) across Infinity markers and algorithms. **b)** Density heatplots of measured (x-axis) versus predicted (y-axis) for 12 Infinity markers sampled across the whole range of performances. Vertical lines indicate the thresholds chosen to define positive expression of the markers. **c)** For each algorithm, distribution of AUC scores for different sizes of the training set. Three markers are individually highlighted. **d)** Runtime for the four algorithms for different sizes of the training set and a fixed size of the imputation set.

Importantly, true staining performance can vary widely across the range of empirical measurements. This can be due to a lack of the marker in the sample (e.g. RBC-lysis during sample preparation precludes detectable expression of the RBC-specific marker Ter-119), sensitivity to tissue preparation method (e.g. CD115 is heavily cleaved by many collagenase enzymes), staining approach (many chemokine receptors require staining at 37°C for robust signal), or simple reagent quality as not all antibodies perform equally well. Thus evaluation of performance must account for this empirical variability as well as algorithm performance. Consistently, performance metrics for Infinity markers fell within a range of AUC from 0.72 to 0.98. We highlight 12 markers sampled across the whole range of performances in Figure 2b. As expected, markers that resulted in high AUC were typically multi-modal and either highly or commonly expressed. Exhaustive manual examination showed that 155 out of 252 phenotypic markers (61.5%, excluding isotype controls and autofluorescence measurements) yielded meaningful imputed signal. For a limited set of seven markers consisting mostly of TCR spectratyping antibodies (e.g. TCR Vβ7, TCRVβ9) the models were unable to accurately impute expression levels despite the presence of a positive cell population (Supplementary Figure 2). This reflects an expected limitation of the approach: the expression of an Infinity marker requires a unique signature in the Backbone data to be accurately imputed. A TCR Vβ7^+^T cell however resemble any other T cell with our Backbone panel. This expected negative result underscores the importance of the Backbone panel design in the Infinity Flow approach. Further, it highlights that future refinement of the choice of Backbone antibodies and tuning methodology to a given sample type may enhance the overall breadth of the assay.

Markers restricted to rare cell populations in general gave lower AUC values, though still well above an AUC of 0.5 (corresponding to a random guess) (Figure 2b). To estimate a minimal number of positive events allowing satisfactory performance when predicting a rarely-expressed marker, we studied the influence of the number of events during training on the performance of these models. We trained each algorithm for each marker on a varying number of training events (from 1,000 to 50,000) and tested accuracy on held-out data. For each algorithm, performance increased with training set size, and plateaued after 10,000 events. Widely expressed markers such as CD48 or CD98 were mostly unaffected by this restriction of the training set size. Performance was however impacted for CD200R3, a marker specifically expressed on basophils which are exceedingly rare in the analyzed dataset (0.82% of the total live single cells). At very low cell numbers (1,000 sampled event per well, or on average 8 basophils and 992 non-basophils during training), XGBoost and SVM still resulted in high AUC (≥ 0.8), suggesting that these methods are able to accurately predict the expression of a marker specific for a rare cell population even when training data is limited (Figure 2c and Supplementary Figure 3).

**Figure 3.**
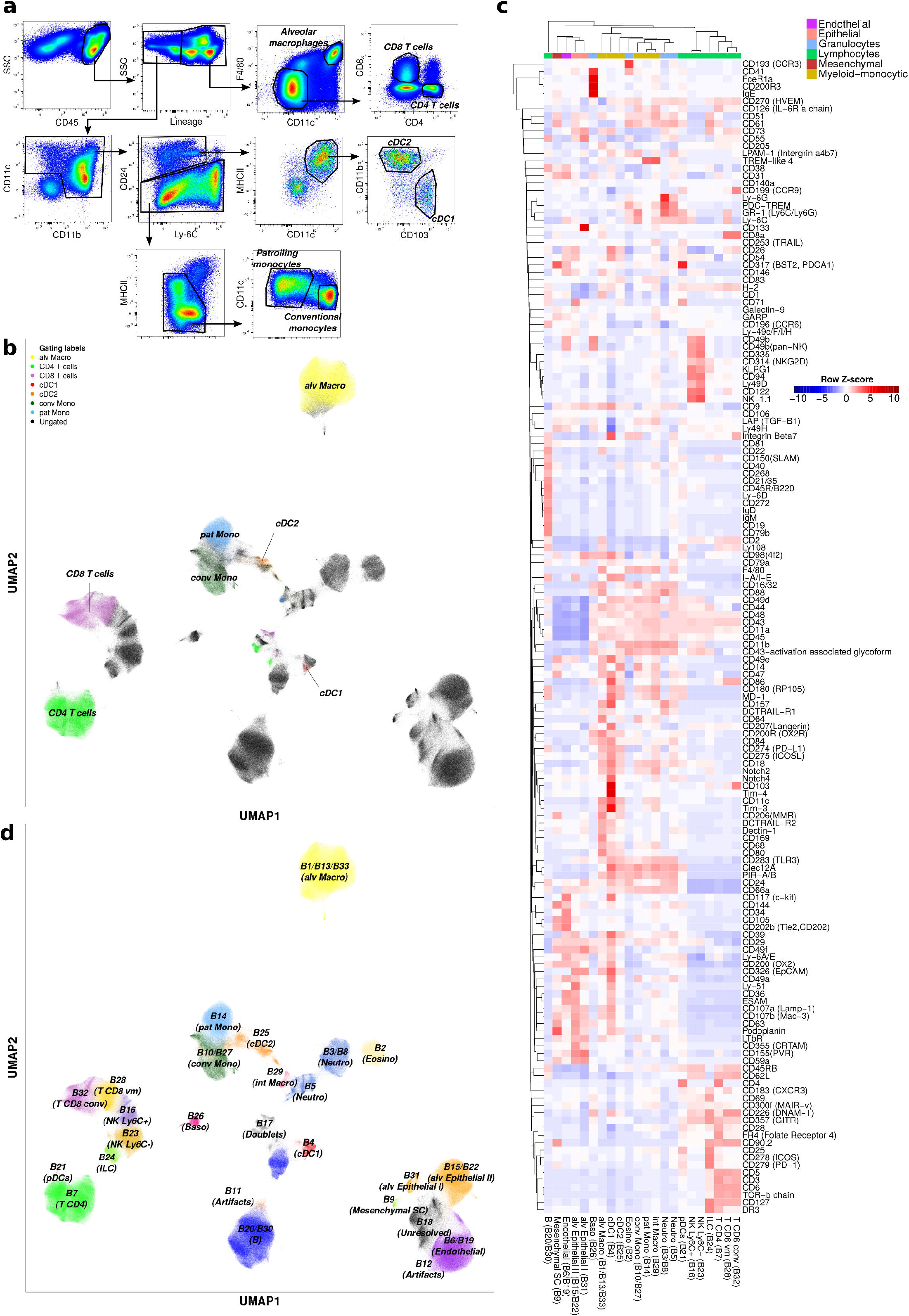
Infinity Flow enables near-exhaustive phenotyping of lung cells. **a)** Manual gating of the Backbone data from cells isolated from the lungs of C57BL/6 mice, and **b)** UMAP dimensionality reduction of the Backbone data, colored by cell phenotypes from manual gating. **c)** Median Infinity Flow predictions of 155 Infinity markers showing staining across cell clusters defined by the Phenograph algorithm on the Backbone data. **d)** Projection of the cell clusters on the UMAP embedding of the Backbone data.

The performance benchmarks described above are performed per well. However, technical variability (batch effects) can further impact prediction accuracy across wells. The goal of the Infinity Flow method is to enable cross-well imputation of marker expression with high accuracy. Thus, to test for the ability of these models to generalize to data independent from a specific well, we compared the cross-well predictions with within-well predictions of the models by rotating-out Backbone markers one at a time and using the remaining Backbone markers to predict the intensity of the left-out marker. The impact of cross-well prediction on performance was limited (median △R^2^ from −0.015 to −0.012 across algorithms, Supplementary Figure 4a). XGBoost was more accurate than other non-linear models for virtually any marker-algorithm pair (Supplementary Figure 4b).

**Figure 4.**
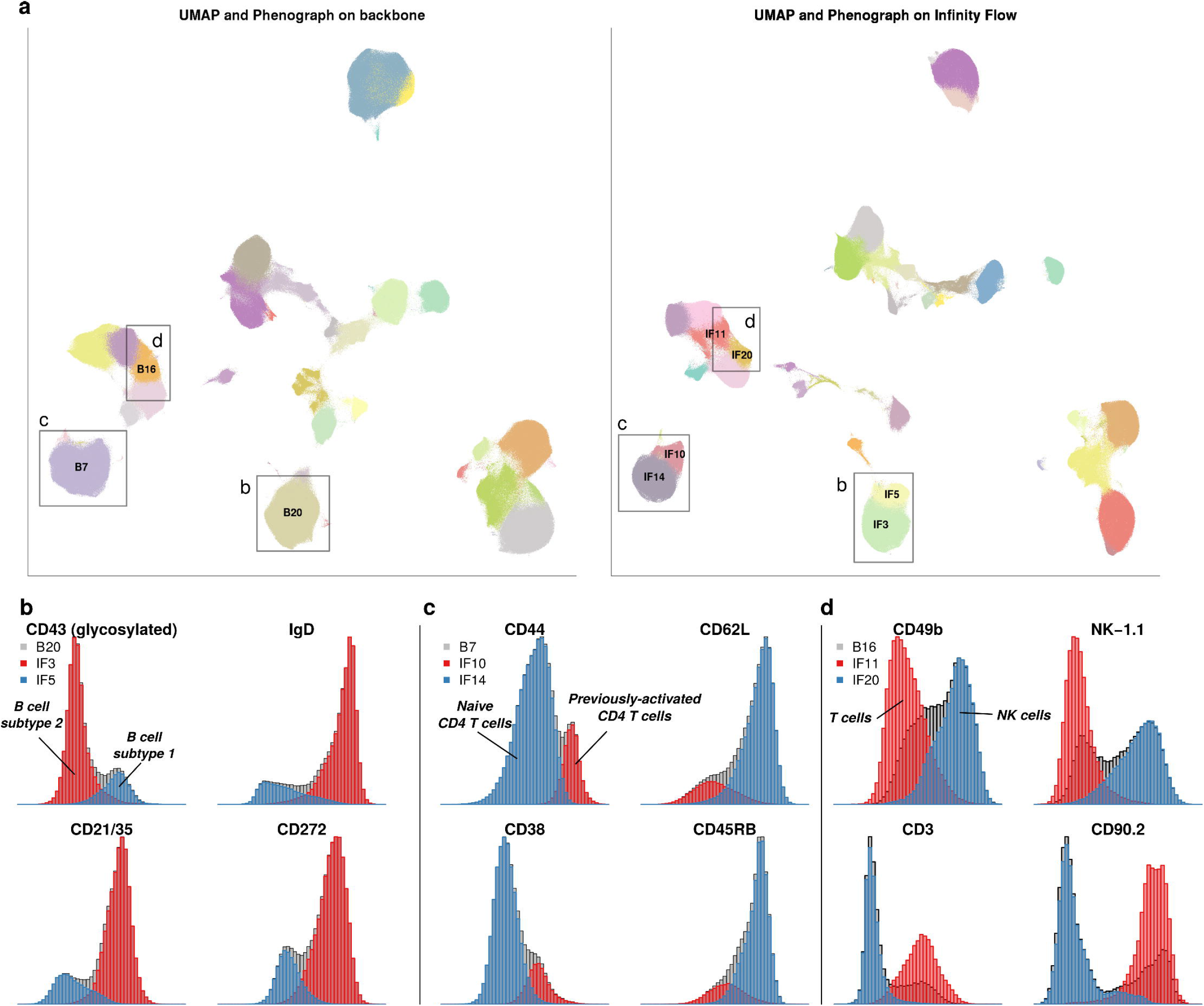
Infinity Flow increases the signal-to-noise ratio of MPC datasets. **a)** Side-by-side Phenograph clustering and UMAP embedding of the Backbone data (left) and the Infinity Flow-augmented dataset (right). **b)** Distribution of markers B cell subtype 1 and B cell subtype 2 in the B cell Backbone cluster B20 (gray) and the two Infinity Flow-augmented B cell clusters IF3 and IF5 **c)** Distribution of markers of naive and previously-activated CD4 T cells in the CD4 T cell Backbone cluster B7 (gray) and the two Infinity Flow-augmented CD4 T cell clusters IF10 and IF14 **d)** Distribution of markers of NK and T cells in the mixed T CD8 and NK cells Backbone cluster B16 (gray) and the two Infinity Flow-augmented clusters IF11 (T cells) and IF14 (NK cells)

Finally, we found that the total runtime of XGBoost was the shortest of all benchmarked methods (Figure 2d). Due to its accuracy at both high and low cell numbers and speed, the XGBoost implementation of gradient boosted trees is the default imputation method offered in the Infinity Flow package. Importantly, all tested non-linear algorithms provided accurate and robust imputed data for most markers, highlighting the robustness of the Infinity Flow approach to the choice of multivariate regression framework used.

### 2.3 Infinity Flow enables cell-level background correction in MPC assays

One of the limitations of flow cytometry data is that the signal on non-expressing cells still follows a non-random pattern, mostly due to autofluorescence and unspecific antibody binding, even with standard Fc-receptors blocking approaches. As such, our derived predictions include a significant and underdispersed contribution from non-specific signal. Using Infinity Flow-imputed data, we were able to enhance the signal of dimly-expressed antibodies by performing cell-wise correction of the non-specific background fluorescence signal: since Infinity Flow jointly imputes the expression of each Infinity marker alongside the “expression” of isotype-matched control antibodies across every cell, we could mathematically model this background signal (Methods). The residuals of these models, akin to per-cell background substracted expression values, were used as background-corrected data (Supplementary Figure 5a). These background-corrected data enabled clear identification of marker-expressing cells for some dimly-expressed markers, such as CD279 (PD-1) (Supplementary Figure 5b), and removed spurious correlation patterns within non-expressing cells (Supplementary Figure 5c, 5d). Of note, this approach could also find applications in conventional flow cytometry, whenever isotype controls are included in the antibody panel, as per cell background correction is easier to interpret than traditional methods based on manual gating.

**Figure 5.**
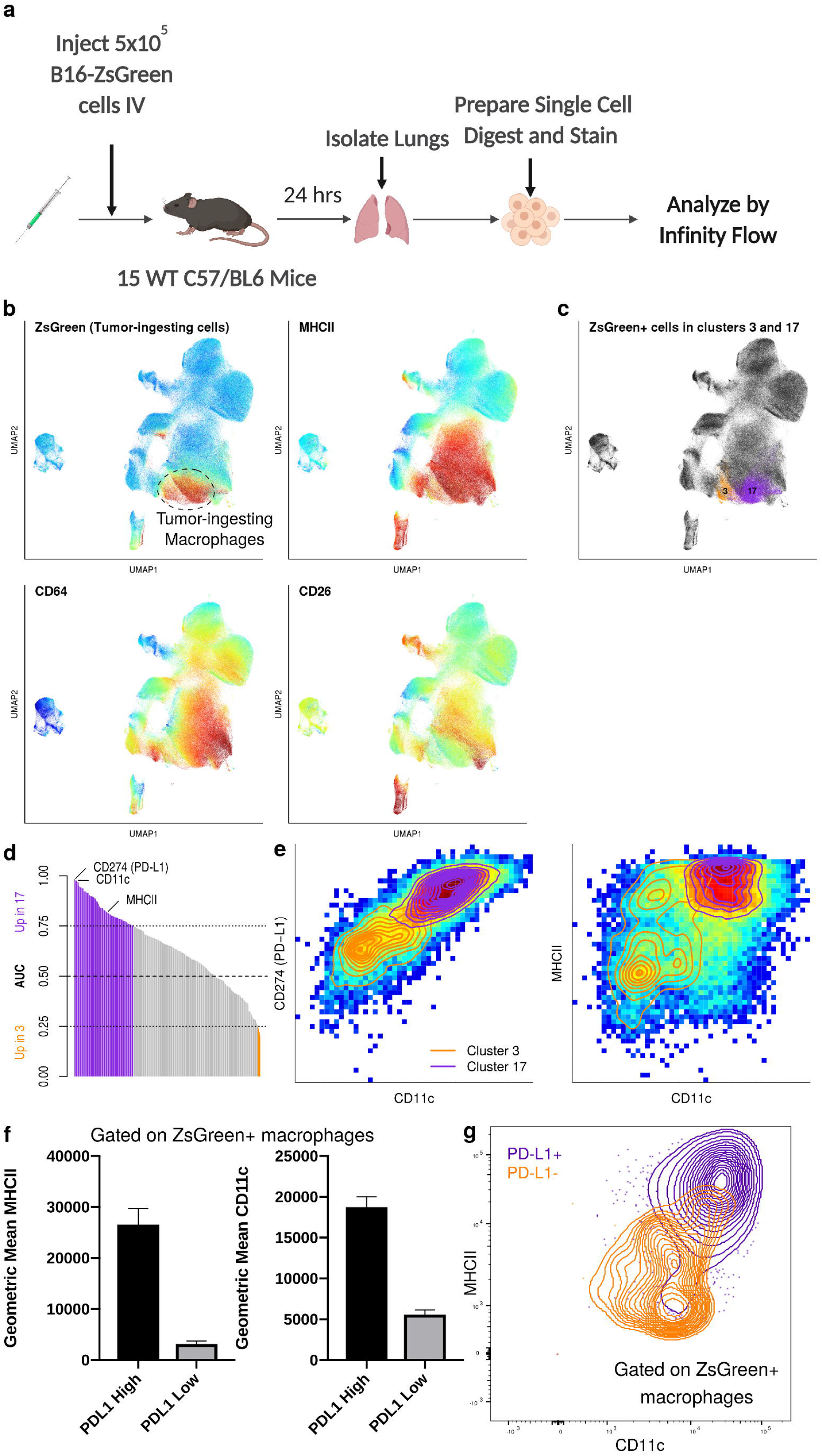
Infinity Flow identifies heterogeneity within tumor-ingesting macrophages during metastasic seeding of the lung. **a)** Outline of the experimental setting. **b)** Color-coded expression of ZsGreen (Backbone) and MHCII, CD64 and CD26 (imputed) on a UMAP embedding of myeloid cells. **c)** ZsGreen^+^ events from two Phenograph clusters of macrophages. **d)** Barplot representing the AUC of every imputed marker for ZsGreen^+^ cells from the two macrophages clusters. **e)** Color-coded densities of single cells for pairs of markers, overlayed with contours of the two macrophages clusters. **f)** Median fluorescence intensities of MHCII and CD11c for PD-L1^+^ and PD-L1^-^ macrophages in an independent validation cytometry experiment. **g)** Contour plot of PD-L1^-^ and PD-L1^+^ macrophages on a CD11c versus MHCII plot.

### 2.4 Infinity Flow enables the comprehensive annotation of the cellular components of the mouse lung

Having established the accuracy and robustness of the Infinity Flow approach, we next sought to utilize it in a real-world experimental context. Our laboratories have extensive experience using flow cytometry to profile the cellular environments of tissues such as lung (Headley et al. 2016). In the past, comprehensive analysis of such a tissue, inclusive of diverse immune and non-immune populations, would have required several multi-parameter flow cytometry panels run in sequence and would still have failed to classify many cell types. Further, comprehensive phenotyping of these cells would have been exceptionally labor-intensive by this standard approach. We thus evaluated whether the Infinity Flow approach could improve upon a conventional multi-color flow cytometry panel and offer new insights into the cellular constituency of complex tissues and disease states.

We first applied Infinity Flow to the complete mouse lung cell dataset (partly shown in Figure 1f). Mouse lung cells were isolated from the unperfused lungs of C57BL/6 mice at the steady state and stained using a 14-color Backbone panel consisting of 11 antibodies in individual channels, 2 multiplexed channels (containing various lineage markers in combination), live/dead stain, and 2 light scatter parameters (Supplementary Table 3). The Infinity Panel consisted of 252 PE-conjugated antibodies and 14 matched isotype and autofluorescence controls (Supplementary Table 4). A total of 28,715,415 live single cell events were acquired. We used 50% of the measured events to train the machine learning models, and output imputed data for w = 266 antibodies across n = 10,000 ×266 = 2, 660,000 single cells distinct from those used for training the models (Supplementary Figure 6).

Our standard gating scheme on the Backbone for a panel of this nature allowed us to account for only 37.7% of live single lung cells (Figure 3a, 3b). UMAP dimensionality reduction (Becht et al. 2019; McInnes, Healy, and Melville 2018) of the Backbone data however revealed additional information pertaining to cell populations within the Backbone measurements and not captured using this standard approach (Figure 3b). Most of the unclassified events were either non-immune (CD45^-^), or positive for the highly-multiplexed lineage channel, comprised of a cocktail of 8 antibodies specific to T cells, B cells, NK cells, neutrophils, alveolar macrophages and eosinophils (Supplementary Table 3). Infinity Flow deconvolved this multiplexed lineage channel into its constituent markers, enabling straightforward and accurate identification of the individual cell types which comprise these events (Supplementary Figure 7). These results highlight the ability of this approach to access and utilize the information contained within these highly multiplexed channels.

To enhance our ability to delineate cell types within the murine lung, we applied Phenograph clustering (Levine et al. 2015) to the Backbone measurements. Centroids of the Infinity Flow-imputed expression intensities across a set of 39 known cell type defining markers across clusters allowed identification of almost every cell cluster (97% of live single cells, 32 out of 33 clusters)(Supplementary Figure 8).

Only a single cluster (cluster 18) that was negative for virtually every lineage marker was left unclassified (Supplementary Figure 8). In addition, combining Infinity Flow-imputed data with Phenograph clustering allowed us to identify and classify rare cell populations of CD200R3^+^ FCϵR1+basophils and of innate lymphoid cells (ILCs). These rare cell populations represented respectively only 0.82% and 1.08% of the total live single cells (Supplementary Figure 9). This analysis thus resulted in a near-exhaustive phenotypic classification of the mouse lung at the single cell level, associated with extensive protein expression profiles for each cell population (Figure 3c). Additionally, this approach enabled classification of several major components of the non-immune constituency of the lung, an unexpected benefit that could be enhanced with Backbone panel design to enable deep simultaneous classification of both immune and non-immune cell-types in the lung (Figure 3c, 3d).

### 2.5 Infinity Flow increases the signal-to-noise ratio of MPC datasets

One of the exciting applications of this method is to better understand and profile the heterogeneity in cell types within a complex tissue. Clustering and non-linear dimensionality reduction algorithms are becoming increasingly common methods to define cell subsets within single cell data. However, these approaches are often unsupervised, and are therefore limited in their ability to distinguish real patterns from noise in data. Since Infinity Flow uses supervised learning methods, we reasoned it could enhance our ability to identify biologically relevant cell populations in Infinity Flow-processed datasets compared to the analysis of the Backbone data alone. To compare the richness of Infinity Flow’s output matrix to the initial Backbone matrix, we re-analyzed the murine lung dataset using UMAP dimensionality reduction and Phenograph clustering (Methods), now including both the Backbone and imputed measurements as input (Figure 4a).

Overall, applying UMAP dimensionality reduction and Phenograph clustering on the Infinity Flow-augmented data lead to more accurate and informative Phenograph-clustering and UMAP-embedding. Notably, it separated B cells into two distinct Phenograph clusters that were well-separated in the UMAP-embedding (Figure 4a, 4b, Supplementary Movie 1). Imputed markers expression suggested that Infinity Flow was able to resolve the recently-described B1 (glycosylated CD43^+^IgD^-^CD21/35^-^CD272^-^) and B2 (glycosylated CD43^-^IgD^+^CD21/35^+^CD272^+^) subsets of B cells in this dataset (Baumgarth 2016; Haas 2015). Similarly, Phenograph clustering of the Infinity Flow-augmented data separated CD4 T cells into two distinct clusters, corresponding to naive (CD44^-^CD38^-^CD62L^+^CD45RB^+^) and previously-activated (CD44^+^CD38^+^CD62L^-^CD45RB^-^) CD4 T cells (Figure 4a, 4c). Phenograph clustering of the Backbone data also created a cluster overlapping the boundary between T cells and NK cells, while Phenograph clustering of the Infinity Flow-augmented data segregated these two cell populations more accurately (Figure 4a, 4d). In addition, the Infinity Flow-augmented UMAP positioned dendritic cells and monocytes clusters in closer proximity, in accordance with their ontogeny (Supplementary Movie 1). These examples illustrate how the single cell resolution and high number of parameters of Infinity Flow’s imputed data allow a more accurate and granular delineation of the cell types present.

### 2.6 Infinity Flow identifies heterogeneity within tumor-ingesting macrophages during metastasic seeding of the lung

A key promise of high-dimensional cellular profiling is the ability to better explore and understand the complexity of cellular phenotypes in disease states. We previously identified a unique axis in the early immune response to pulmonary metastasis wherein lung macrophages encounter and ingest tumor material in the metastatic niche (Headley et al. 2016). This earlier analysis was however limited to a relatively narrow immune-profiling panel (Supplementary Table 3) which by itself lacked the capacity to finely delineate myeloid subsets within this early disease state. In order to assess Infinity Flow’s ability to profile and detect subtly distinct cell populations within the complex environment of early lung metastasis, we stained cells from the lungs of mice implanted with metastatic ZsGreen-transfected B16 melanoma cells 24 hours prior. In this system the B16 melanoma cells express the green fluorescent protein, ZsGreen, enabling identification of phagocytes that have ingested tumor cells (Figure 5a). Our prior studies identified a population of macrophages which critically support metastatic growth. A portion of these macrophages ingest tumor material very early in the process of metastatic seeding. Consistently with our prior analysis, the majority of tumor ingesting (ZsGreen^+^) immune cells within the myeloid compartment phenotypically fit the profile of macrophages, expressing high levels of CD64, low levels of CD26, and in this context high levels of MHCII (Figure 5b). Intriguingly, Phenograph clustering of Infinity Flow’s output revealed two distinct clusters of ZsGreen^+^ macrophages (clusters *3*and *17,* Figure 5c and Supplementary Figure 10). This heterogeneity was not appreciated in our previous analysis using conventional gating (Headley et al. 2016). We next sought markers distinguishing these two clusters of tumor-ingesting macrophages. A variety of markers showed distinct expression patterns between the two clusters and we selected three among the ones that best separated the two clusters (Figure 5d). We found that a combination of CD11c, MHC-II, and PD-L1 effectively defined each cluster, with cluster *3* being low and cluster *17* high for each of these markers (Figure 5e). To validate these findings, we replicated this same stain via conventional flow cytometry in a separate cohort of mice. We were able to confirm the presence of both PD-L1^high^ and PD-L1^low^subsets of metastasis-ingesting macrophages in all mice, with a phenotype consistent with those observed in the Infinity Flow dataset (Figure 5f, 5g). Further exploration is required to ascertain the functional relevance of these two distinct populations of metastasis ingesting macrophages, but these validated findings demonstrate the potential for Infinity Flow in interrogating the cellular constituencies of disease states and deriving new experimental hypotheses.

## 3 Discussion

In this article, we described Infinity Flow, a combined experimental and computational workflow enabling the analysis of the expression and co-expression patterns of hundreds of surface proteins across millions of single cells. The Infinity Flow computational pipeline is available as an open-source R package, and can run on standard FCS files generated from massively-parallel cytometry experiments. Others have published large scale antibody screens using MPC experiments, but the analysis of these datasets has been limited to manual or automated analysis of the Backbone data (Amir et al. 2019; Collier et al. 2017; Graessel et al. 2015; Koh et al. 2016; Uezumi et al. 2016; Kalina et al. 2019). In contrast, Infinity Flow uses supervised machine learning (specifically multivariate non-linear regression algorithms) to aggregate expression data from all the well-specific FCS files into a single dense matrix of imputed expression. To achieve this, we leverage the fact that the Backbone markers are measured for every cell in every well and thus allow detection of non-linear correlations between expression levels of the Backbone markers and the Infinity marker for that well. This information is then utilized to model and impute expression values for all markers on every cell in the study. Our study suggests that non-linear machine learning models vastly outperform linear models for this task.

By imputing the expression levels of hundreds of proteins across millions of single cells, Infinity Flow enables broad phenotypic characterization of many cell subsets at the level of a whole organ. The depth of this characterization is limited by the complexity of the Backbone antibody panel. For this approach to work, it is essential that Backbone markers do in fact inform upon the expression of the Infinity marker. We notably highlighted that the Backbone panel we used to analyze the mouse lung at steady-state was unable to accurately predict expression patterns of specific TCR antibodies (e.g. Vb7, Va3.2) despite clear staining of discrete T cell sub-populations in the Infinity antibodies measurements (Supplementary Figure 2). Extra Backbone antibodies could allow more depth by using antibodies specifically targeting rare cell subsets (e.g. to resolve *T_γδ_* cells from T_αβ_ cells). To produce the datasets analyzed in this article, we used a relatively modest LSRFortessa cytometer. State-of-the-art instruments such as spectral flow cytometers should allow the resolution of an increased number of cell subsets while still increasing the overall number of parameters that can be analyzed many-fold over what is currently possible on even the most advanced of these instruments. Our results highlight the ability of Infinity Flow to deconvolve highly multiplexed acquisition channels. This was notably made possible by designing a multiplexed channel with distinct modes within positive cells. By optimizing the multiplexing of more channels, one could likely achieve high cellular resolution while using only a limited number of channels, and therefore successfully use the Infinity Flow pipeline on even the most modest of cytometers. The design of the backbone panel, while critical, does not differ much from the design of a traditional antibody panel: the goal remains to maximize the cellular resolution, with the added constraint that spectral overlap with the variable channel should be minimized. The broad characterization of the mouse lung achieved herein however highlights that Infinity Flow is well-suited to deeply characterize the cellular composition of complex tissues at the protein-level. In this article we only analyzed single-sample datasets. Conventional antibody-based barcoding of multiple samples mixed in equal cellular proportions should however enable multi-samples Infinity Flow. An alternative approach to MPC is the use of multiple variable antibodies per well, as recently performed by Glass *et al.* using mass-cytometry (Glass et al. 2020). The computational pipeline introduced herein straightforwardly extends to this type of dataset. This method and our Infinity Flow method represent complementary approaches to high-dimensional flow cytometric analysis of tissues allowing researchers to select the method of choice based on their access to technology. The Glass *et al.* approach is suited best to mass cytometry-based technology, as multiple complex panels need to be simultaneously optimized. In mass cytometry, spectral overlap across channels is limited. This is not the case for flow cytometry, which largely complicates the simultaneous measurement of multiple variable antibodies. Infinity Flow requires a single backbone panel which in our experience can be used across an array of tissues and contexts. Further, the approaches differ in throughput as Infinity Flow can easily allow sampling on the order of 10s of millions of cells while mass cytometry-based approaches are more limited in cell throughput, although both methods offer higher throughput than sequencing based approaches.

The critical importance of the Backbone panel for imputation highlights that any pattern captured by Infinity Flow has an underlying signature in the Backbone data. As the Backbone is an information bottleneck in MPC assays, one could intuitively expect that performing dimensionality reduction or clustering on the Infinity Flow-augmented data matrix would yield similar results to performing these analyses on the Backbone data directly. Importantly, we found that the augmented data matrix enhanced our ability to identify cellular heterogeneity within individual cell clusters. We believe a key element enabling this is the use of supervised machine learning models that are capable of highlighting subtle patterns in the Backbone data as relevant for the prediction of a given Infinity antibody, a process comparable to feature engineering for machine learning. These results highlight the complexity of inferences which can be derived from multicolor flow cytometry data when treated as continuous, with interrogation of marker-marker relationships in an unbiased and non-linear fashion.

Compared to other high-dimensional single-cell protein expression approaches such as oligonucleotide-tagged antibodies (Stoeckius et al. 2017; Shahi et al. 2017) or mass-cytometry (Bendall et al. 2011), Infinity Flow is affordable, allows many cells to be profiled thanks to the high cellular throughput of conventional flow cytometry, and does not require dedicated equipment or expertise other than those used for conventional flow cytometry. The increased cellular throughput notably allows profiling of both abundant and rare cell populations in a single experiment without pre-enrichment. Issues that prevent high parameterization with other methods, such as fluorescence overlap and availability of unique fluorophores (conventional flow cytometry), availability of unique metals (mass cytometry) or available sequencing space (CITE-seq or Abseq) are therefore not a hindrance to the Infinity Flow method. There is no theoretical limit to the number of surface proteins that can be assessed. All staining of Infinity markers is done in parallel wells, using antibodies conjugated to the same common fluorophore (PE or APC, in general). The primary factors restricting the total number of parameters that can be stained are the availability of high-quality fluorescently conjugated-antibodies, and enough single cells to divide the initial sample across as many wells as there are Infinity markers. Generally, between 10^5^ and 10^6^ cells are required per well to account for cell loss during staining while still enabling detection of rare populations. We previously demonstrated the ability for Infinity Flow to to profile pre-enriched rare human circulating cDC2s (Dutertre et al. 2019), and herein show that it can inform upon the cellular composition and phenotypes of a whole organ, including rare cell populations, without pre-enrichment. As Infinity Flow is highly scalable, samples with a relatively small number of cells can still be interrogated by limiting the number of Infinity markers to a smaller subset of interest. Conversely, the studies presented here make use of commercially available kits with several hundred pre-conjugated antibodies. These kits can however be readily augmented by adding additional wells with a custom set of Infinity antibodies best suited to the specific scientific question.

Given the wealth of information provided by the Infinity Flow pipeline, its affordability and compatibility with standard flow cytometers, we anticipate it will be widely adopted.

## 4 Methods

### 4.1 Data generation

#### 4.1.1 Preparation of single cell lung suspensions

For all experiments single cell digests were prepared from lung as previously described (Headley et al. 2016). Briefly, lungs were collected from 15 8 week old male C57BL/6 mice following euthanasia by overdose with 2.5% Avertin. For steady-state lung analysis, mice were previously unmanipulated. For analysis of metastasis bearing lungs - all mice were injected 24 hours prior with 5 ×10^5^ZsGreen-expressing B16-F10 melanoma tumor cells via the tail vein, as previously described (Headley et al. 2016). Following harvest, lungs were suspended in 5 mL DMEM (GIBCO) with a combination of digestive enzymes (0.26 Units/mL-1 LiberaseTM (Sigma) and 0.25 mg/ml DNaseI (Sigma)). Samples were placed in C-Tubes (Miltenyi) and briefly processed with a GentleMACS Octomacs Tissue Dissociator (Miltenyi) using the standard Lung processing protocol built into the instrument. Samples were then incubated at 37°C for 30 min with shaking at 150rpm and processed a second time via GentleMACS. Samples were filtered through a 100μm cell strainer (Fisher), centrifuged at 300g for 5 min and resuspended in 3 mL of 1X ebioscience Red Blood Cell Lysis Buffer (ThermoFisher) for 5 min at 37°C. Lysis buffer was quenched with 10mls of FACS buffer (PBS + 2% FCS) and centrifuged at 300g for 5 min. Samples were resuspended in PBS and then filtered through a 40 μm cell strainer (Fisher) all samples were then pooled and cell concentration was adjusted to 20 ×10^6^ cells/mL for staining.

#### 4.1.2 Massively parallel flow cytometry staining

All MPC experiments in this protocol were performed with Biolegend LEGENDScreen Mouse PE Kit (Biolegend) and manufacturer recommendations for LEGENDScreen Plate preparation and staining were performed as indicated by the manufacturer, unless noted otherwise below. In brief, single cell suspension of lung cells were washed and resuspended at 20 ×10^6^ cells/mL in PBS + Zombie NIR Live/Dead stain (Biolegend) at 2μL/mL and incubated for 20 min at room temperature. Following staining, a 10-fold volume of Cell Staining Buffer (Biolegend) was added to neutralize any unbound dye and cells were centrifuged at 300g for 5 minutes. Cells were resuspended at 20 ×10^6^ cells/mL in Cell Staining Buffer and non-specific staining was then blocked by addition of 2μg/mL anti-CD16/32 (mouse TrueStain FcX Biolegend), and 2% rat serum (Invitrogen) and 2% Armenian hamster serum (Innovative Research) followed by 15min incubation at 4^°^C. Cells were then washed and resuspended in Cell Staining Buffer and a master-mix of the indicated Backbone antibody panel (Supplemental Table 1) at 20 ×10^6^ cells/mL and incubated 30min at 4°C. Following Backbone staining cells were washed twice and resuspended at 20 ×10^6^ cells/mL in Cell Staining Buffer and 75μl were added to each of well of the LEGENDScreen plates. Staining and fixation were performed exactly as per manufacturer directions. Following staining, each independent well was filtered using Acroprep 40μm 96-well filter plates (Pall Laboratories) and the final volume adjusted to 200μL of Cell Staining Buffer. For tumor metastasis experiments, cell fixation was not performed in order to maximally preserve ZsGreen fluorescence. Note: While these experiments were performed using Biolegend LEGENDScreen kits (Biolegend), BD Lyoplate Mouse Cell Surface Screening Panels (BD Biosciences) have also been utilized for MPC staining with similar results. However, the Lyoplate method utilizes unlabeled primary antibodies and secondary staining with biotinylated-anti IgG secondary antibodies followed by detection with streptavidin-A647 Backbone panel staining, thus Backbone panel staining has to be performed in well subsequent to staining with the Infinity panel.

#### 4.1.3 Flow cytometric data collection and data preprocessing

Samples for these experiments were collected on a BD Fortessa X-20 cytometer using the BD HTS system to sample from each well independently. Compensation was performed using ebioscience UltraComp beads (ThermoFisher) stained with Backbone panel antibodies and Ly-6G (Clone 1A8) conjugated to PE to ensure a very bright signal in the PE channel. Compensation for Zombie NIR was performed using BD Amine Reactive Beads (BD Biosciences) stained as per manufacturer protocols. 50% of the total volume of cells were run (100μL) enabling rerunning of any sample where clogging or other technical issues prevented clean collection of data. Following data collection, FCS files were individually examined in FlowJo (Treestar Inc) for quality control. Each fluorescence channel was plotted against time to assess technical artifacts due to cytometry clogging - these individual samples were recollected if necessary. Fluorescence compensation was also checked for accuracy and adjusted if necessary. Singlet and live cell gating was then performed consistently for each FCS file and this preprocessed data (well compensated, live, singlets) were exported into individual FCS files for downstream input into the Infinity Flow pipeline as described below.

### 4.2 Computational analyses

#### 4.2.1 The *infinityFlow R package*

Infinity Flow is implemented as an R package, *infinityFlow* (https://www.github.com/ebecht/infinityFlow). A flow chart summarizing the key steps of the pipeline is shown is Supplementary Figure 11. The input data consists of a set of FCS file, one per MPC well. These files can be uniformly downsampled in the event that computational resources are limited. The input data is then transformed using logicle transformations (Parks, Roederer, and Moore 2006) with parameters estimated as in the flowCore R package (Hahne et al. 2009). In order to harmonize data across input wells, each Backbone channel is linearly-transformed to have 0-mean and unit variance (Z-score) within each well. Non-linear multivariate regression models are then trained and used to impute Infinity antibodies’ expression intensities as described below. UMAP non-linear dimensionality reduction is run on the concatenated Backbone data matrix. The output matrix contains the Backbone measurements, imputed Infinity antibodies expression values, UMAP coordinates and well identifiers for each event. Another similar output matrix is produced using background-corrected signal, as described below. For each of these output matrices, a concatenated FCS file is output, as well as one FCS file per input well. Finally, a UMAP plot color-coded by the measured or imputed expression of each Backbone or Infinity marker is produced, similarly to Supplementary Figure 6.

#### 4.2.2 Regression models

The *InfinityFlow* R package currently supports four classes of non-linear multivariate regression models. These four methods are Support Vector Machines (SVM) (Chang and Lin 2011) implemented in the *e1071* R library, Gradient Boosted Trees implemented in the *xgboost* R library (Chen and Guestrin 2016), neural networks using *the* tensorflow and *keras* R packages (Chollet and Allaire 2018), as well as polynomial models of arbitrary degrees, with or without L1-regularization (Tibshirani 1996), implemented in the *glmnet* R library. One regression model per algorithm and Infinity antibody is fit. The data is first randomly split into a training set (50% of the events) and a test set to evaluate performance (50% of the events). Finally, the size of the output is chosen by the user, and drawn from the test set for downstream analyses. For all models, the target variable is the intensity of the Infinity antibody, while the predictor variables are intensities of Backbone antibodies.

#### 4.2.3 Regression models’ hyperparameters

The regression models’ hyperparameters can be chosen by the user. For convenience, we provide default values for each regression model that provided the best performance on a subset of the data on which we benchmarked these hyperparameters. These hyperparameter values were used throughout the manuscript. For SVM, these arguments’ values are nu-regression with a radial basis function kernel and nu = 0.5. For XGBoost, we used nrounds = 1500 and eta = 0.03. For neural networks, we used a fully-connected neural network with one hidden layer of the same size as the input layer, rectified linear-unit as activation function for nodes in the hidden and input layers, a minibatch size of 128, stochastic gradient descent for optimization with a learning-rate of 0.005, mean-squared error as a loss function, 20% of the training events as a validation set, and up to 1,000 epochs during training, with an early-termination criteria of 20 rounds without reaching a new minimum of the loss function in the validation set. Adding additional hidden layers or using tanh as activation function did not substantially impact performance of the NN models, while training with more examples to accommodate events used for performance evaluation and early stopping had a marginal positive impact on performance that remained lower than the one of XGBoost models (Supplementary Figure 12). For LASSO models, we used either 1st or 3rd degree polynomials, with an amount of L1-penalty automatically chosen using 10-fold cross-validation.

#### 4.2.4 Isotype-specific background staining correction

The massively-parallel flow cytometry kit used herein contains isotype control antibodies for each of the antibody isotypes present. Since Infinity Flow imputes every Infinity antibody, it produces imputed expression levels both for a given Infinity antibody *A* and its corresponding isotype control *I*. To correct for unspecific staining, we fit the following linear model: I =*β_0_*+*Aβi*+ϵ where the residuals e are used as background-corrected intensities.

#### 4.2.5 Dimensionality reduction

For dimensionality reduction, we used the UMAP algorithm (McInnes, Healy, and Melville 2018) using its R implementation in the *uwot* package. UMAP was run with parameters n_neighbors = 15, min_dist = 0.2, metric =“euclidean”, n_epochs =2000. To run UMAP on imputed data, we first removed every events whose PE-intensity was higher than 1/32 of the cytometer’s manufacturer reported maximum linear range, as non-linear effects were causing compensation issues for these events (Supplementary Figure 13), and applied background correction to the imputed intensities. To plot color-coded markers’ intensities on UMAP embeddings, we truncated the imputed intensities data vectors to their lower and upper 5/1000 quantiles.

#### 4.2.6 Performance metrics

To assess the performance of the regression models, we first binarized each Infinity antibody’s PE intensity using manual gating. We used this binary vector and the continuous vector of imputed expression intensities to compute the multiclass area under the receiver operating characteristic curve (AUC) (Hand and Till 2001). To assess the ability of the different models to make predictions that generalize across wells, we iteratively selected one Backbone markers and used the remaining Backbone markers to predict it, using Pearson’s r correlation coefficient as a performance summary. We used paired one-sided Student’s t tests to compare the performance of XGBoost to the other non-linear algorithms benchmarked. To evaluate each model’s runtime, we used wall-clock timing and 24 Intel Gold 6146 cores. To evaluate whether the models overfitted, we examined the average R^2^ coefficient between the predicted and measured data. This analysis revealed similar performance between the training and test sets for neural networks and LASSO models, while the performance of XGBoost and SVM models decreased on the training set. An a posteriori analysis of the average R^2^ coefficient for increasingly many rounds of boosting showed that the performance of the models plateaued rather than decreased on the test set, suggesting that this gap in performance did not impact the practical utility of the XGBoost models (Supplementary Figure 14).

#### 4.2.7 Clustering

Clustering was performed using the Phenograph clustering method (Levine et al. 2015), with parameters k=50. We used the *Rphenograph* implementation of the Phenograph method, using the HNSW library to perform the approximate nearest-neighbor search.

## 5 Data availability

The input, raw predictions and background-corrected predictions for the murine lung dataset at steady state are available at https://flowrepository.org/id/FR-FCM-Z2LP

## 6 Software availability

The development version of the Infinity Flow software is available at https://github.com/ebecht/infinityFlow, and is submitted to the Bioconductor repository.

## 8 Supplementary Figures legend

**8.1 Supplementary Figure 1**

Dotplots of per-well AUC for each pair of non-linear regression models. The dashed line highlights the identity line.

**8.2 Supplementary Figure 2**

Measured versus imputed expression for seven markers for which the Backbone was too limited to inform positive expression. Vertical lines indicate the thresholds chosen to define positive expression of the markers.

**8.3 Supplementary Figure 3**

Measurements (left) and imputed expression (right) for the basophil-specific CD200R3 marker. Imputed expression values are shown for the four non-linear regression algorithms and varying number of cells used for training the models. The zoomed area correspond to the cluster of basophils.

**8.4 Supplementary Figure 4**

**a)** Distribution of change in variance explained (R^2^) when applying a model fit on a given well to other wells, when predicting Backbone markers. **b)** *R* when predicting a given Backbone markers from the remaining Backbone marker. Bar heights represent the *R^2^* averaged across wells, and the bars the corresponding standard deviations. Asterisks represent p-values lower than 0.05 for XGBoost against each other method (paired one-sided Student t-tests).

**8.5 Supplementary Figure 5**

**a)** Imputed expression of the CD4 marker (y-axis) versus imputed signal for the corresponding isotype control antibody. The dashed line corresponds to the ordinary least square fit. 100 random datapoints are highlighted with black hollow circles. The vertical bar represent the corresponding residuals, their lengths being defining the background-corrected imputed CD4 signal. **b)** biplot of raw imputed CD4 and CD8 co-expression, showing either raw (left) or background-corrected (right) values. **c)** biplot of raw imputed

CD4 and PD-1 (a dimly expressed marker) co-expression, showing either raw (left) or background-corrected (right) values. **d)** UMAP embedding of raw predicted values (left) and background-corrected imputed values (right).

**8.6 Supplementary Figure 6**

UMAP embedding color-coded by imputed intensity of each antibody of the Infinity Panel. Phenograph clustering and annotated cell phenotypes are shown in the bottom right corner.

**8.7 Supplementary Figure 7**

UMAP embedding color-coded by measured (for the lineage channel) or imputed (all the other markers) intensities of each antibody used within the Lineage antibody cocktail.

**8.8 Supplementary Figure 8**

Heatmap median values for a curated set of informative markers across Phenograph clusters.

**8.9 Supplementary Figure 9**

Barplot of cell phenotype frequencies in the mouse lung dataset, using Phenograph clusters to define phenotypes.

**8.10 Supplementary Figure 10**

Color-coded Phenograph clustering on a UMAP embedding of myeloid cells in the early lung metastasis dataset.

**8.11 Supplementary Figure 11**

Flowchart describing the Infinity Flow computational pipeline.

**8.12 Supplementary Figure 12**

Performance metrics for different depth, activation functions and training set sizes for neural network models.

**8.13 Supplementary Figure 13**

UMAP embedding color-coded by intensity of the PE antibody measurement (left), imputed values for an antibody creating non-linear compensation artefacts in the Backbone data (middle), and in red events above 1/32 of the manufacturer-documented maximum linear measurement range (right).

**8.14 Supplementary Figure 14**

**a)** Average R^2^ coefficient for the measured versus imputed data on the training and test set for each algorithm used. **b)** Number of boosting steps versus average performance on the training and test set for XGBoost models.

**9 Supplementary Tables legend**

**9.1 Supplementary Table 1**

Typical costs associated with an Infinity Flow experiment

**9.2 Supplementary Table 2**

AUC deciles for each evaluated regression model in the mouse lung dataset.

**9.3 Supplementary Table 3**

Backbone antibody panels for the mouse lung and early lung metastasis datasets.

**9.4 Supplementary Table 4**

Infinity antibody panel used herein.

**10 Supplementary Movie legend**

**Left)** Linear interpolation between the UMAP embedding of the Backbone dataset to the UMAP embedding of the Infinity Flow-augmented dataset. Colors indicate Phenograph clustering. **Right)** Markers of B cells and B cell subtypes 1 and 2, blue indicate low expression, grey intermediate expression and red high expression.

## 11 Abbreviations

AUC: Area under the receiver operating characteristic curve
CD: Cluster of differentiation
ILC: Innate lymphoid cells
MPC: Massively-parallel cytometry
PE: Phycoerythrin

## 12 ACKNOWLEDGEMENTS

We thank Matthew Krummel, Immanuel Kwok, Lai Guan Ng, Robert Balderas, Patrick Oconnell, Rasha Msallam, Jonas Øgaard, Maximilien Evrard, Laura McKay and Robert Amezquita for sharing MPC datasets and/or feedbacks on our software or results. We thank Evan Greene, Nathan Yee and Adam Wojno for reading and critically commenting on the final manuscript. We thank Andrew Berger and the Fred Hutch Flow Cytometry Core as well as the UCSF Flow Cytometry Core for guidance and provision of instrumentation for the experiments described herein. Most of the computational work presented herein has been performed thanks to the resources of the Scientific Computing department of the Fred Hutchinson Cancer Research center. EB and RG were funded by grant NIH 5U19AI128914.

